# Gold Electron Microscopy Grids with Anisotropic Foil Geometry Enable On-Grid Contact Guidance

**DOI:** 10.64898/2026.06.18.733150

**Authors:** Amit Avrahami, Noa Ben Asher, Ran Zalk, Leeya Engel

## Abstract

All-gold electron microscopy (EM) grids reduce beam-induced motion relative to conventional holey carbon supports and provide biocompatible substrates for cellular cryo-EM. However, placing customizable all-gold grid fabrication in the hands of researchers requires accessible processes based on standard microfabrication tools. We report a wafer-scale process using microfabrication techniques available in most academic cleanrooms such as lift-off metallization, electroplating, and sacrificial layer release to fabricate 594 all-gold grids per 4-inch wafer without individual grid handling. A numerical electroplating model provides a quantitative framework to relate gold deposition, grid-bar thickness, and tilt-compatible grid geometry. We show that oval 2 µm × 6 µm foil holes bias on-grid actin organization by substrate geometry alone, without chemical micropatterning. The EM grids supported a 2.15 Å apoferritin single-particle reconstruction on a 200 kV cryo-TEM and are compatible with protein micropatterning and cell culture. This platform establishes an accessible route to programmable, application-specific all-gold cryo-EM supports that couple high-resolution structural imaging with engineered control of cellular organization.

---

Cryo-electron microscopy (cryo-EM) has emerged as a transformative tool in structural biology, enabling molecular-scale visualization of cryogenically preserved protein complexes. A rapidly growing subfield, cryo-electron tomography (cryo-ET), enables three-dimensional visualization of macromolecular structures within frozen-hydrated cells.^1–4^ Both single-particle analysis (SPA) cryo-EM and cryo-ET rely on specialized specimen supports known as electron microscopy (EM) grids. EM grids consist of a conductive structural mesh, 3.05 mm in diameter, overlaid with a thin, typically porous foil.^5^ Whereas SPA studies commonly employ copper meshes, cryo-ET studies involving live-cell culture require biocompatible support materials such as gold or silicon.

For several decades, the state-of-the-art EM support foil consisted of a holey amorphous carbon thin film (atop the metal) mesh with precisely defined hole size and spacing, as exemplified by Quantifoil supports.^6^ Russo and colleagues demonstrated that replacing holey carbon foils with gold foils substantially reduces beam-induced motion, thereby improving cryo-EM image quality. ^7–9^ More recently, Naydenova and Russo introduced an integrated wafer-scale process for batch manufacturing all-gold EM grids with submicrometer holes optimized for SPA.^10^ This work established the feasibility of integrated all-gold EM grid manufacturing and marked an advance in grid manufacturing as previously, the mesh and foil needed to be manufactured separately and subsequently assembled. However, although that work established an important integrated route to all-gold grid manufacturing, reproducing the full workflow in many academic settings can be challenging because key steps, particularly Talbot lithography and cryogenic gold evaporation, require specialized capabilities that are not routinely available in shared university cleanrooms. A broadly accessible process based on standard cleanroom techniques, such as photolithography, lift-off metallization, electroplating, and sacrificial-layer release, would make all-gold grid fabrication more readily reproducible across university nanofabrication facilities and would enable rapid customization of foil and mesh architectures.

For cryo-ET studies of cellular ultrastructure, guiding cells to adopt physiologically relevant morphologies is advantageous because cell shape strongly influences intracellular organization and cytoskeletal architecture.^11–13^ One strategy for directing cell orientation and migration is contact guidance, in which cells respond to physical substrate features or topographical cues.^14–16^ We recently applied a contact-guidance approach to cryo-ET using electrospun fibers to promote cellular elongation and alignment on EM grids.^17^ More commonly, cell shape and positioning on EM grids are controlled using chemical micropatterning approaches for downstream cryo-ET studies.^18–21^ For example, elongated rectangular micropatterns can guide cardiomyocytes toward the elongated morphologies that better approximate their native *in vivo* organization.^22,23^ Despite their utility, these strategies involve processing steps that are additional to and separate from the grid fabrication itself, and require dedicated instrumentation such as maskless UV-exposure systems or electrospinning setups. An approach that encodes a cell-guiding function directly in the grid’s foil architecture at the fabrication stage would eliminate separate surface-chemistry steps and allow the guide cue to be defined by foil design alone.

Here, we present a fully integrated and scalable process for batch fabricating all-gold EM grids using standard semiconductor manufacturing equipment commonly available in university cleanroom facilities. To tune grid-bar thickness and tilt access, we developed a numerical electroplating framework that relates current density, deposition geometry, and the time-dependent evolution of gold deposition during fabrication. Beyond scalable manufacturing, we leveraged the versatility of the process to engineer anisotropic foil geometries that provide intrinsic contact-guidance cues directly at the EM grid surface. Oval foil holes present anisotropic contact-edge geometry to cells at the foil–cell interface, providing directional topographic cues that we hypothesized would direct cytoskeletal reorganization along the hole’s major axis. ^14–16^ We show that replacing circular holes with oval holes is sufficient to bias on-grid actin orientation without altering surface chemistry. Together, these capabilities establish a customizable platform for tailoring mesh and foil architectures to create application-specific supports cryo-EM studies.

## Results and Discussion

### Wafer-scale fabrication of all-gold EM grids

We designed an integrated wafer-scale process for fabricating all-gold EM grids using standard microfabrication methods (Figures 1 and 2). The workflow defines the thin holey foil first, builds the structural grid bars by electroplating, and then releases the completed all-gold grids from the sacrificial wafer. A titanium/gold holey foil is first defined on a fused-silica 4-inch wafer by lift-off. A second lithography step defines the grid bars and perimeter in thick photoresist, after which gold electroplating builds the structural mesh. Finally, hydrofluoric acid removes the fused silica and titanium sacrificial layers, releasing 594 individual all-gold grids. A single fabrication cycle requires approximately four days, and parallel wafer processing enables scalable throughput. Because the foil-hole geometry is defined lithographically and transferred to the metal layer by lift-off, changing the photomask is sufficient to alter the foil-hole geometry without changing the rest of the fabrication workflow. We used this design flexibility to fabricate the oval-hole supports described below.

**Figure 1:**
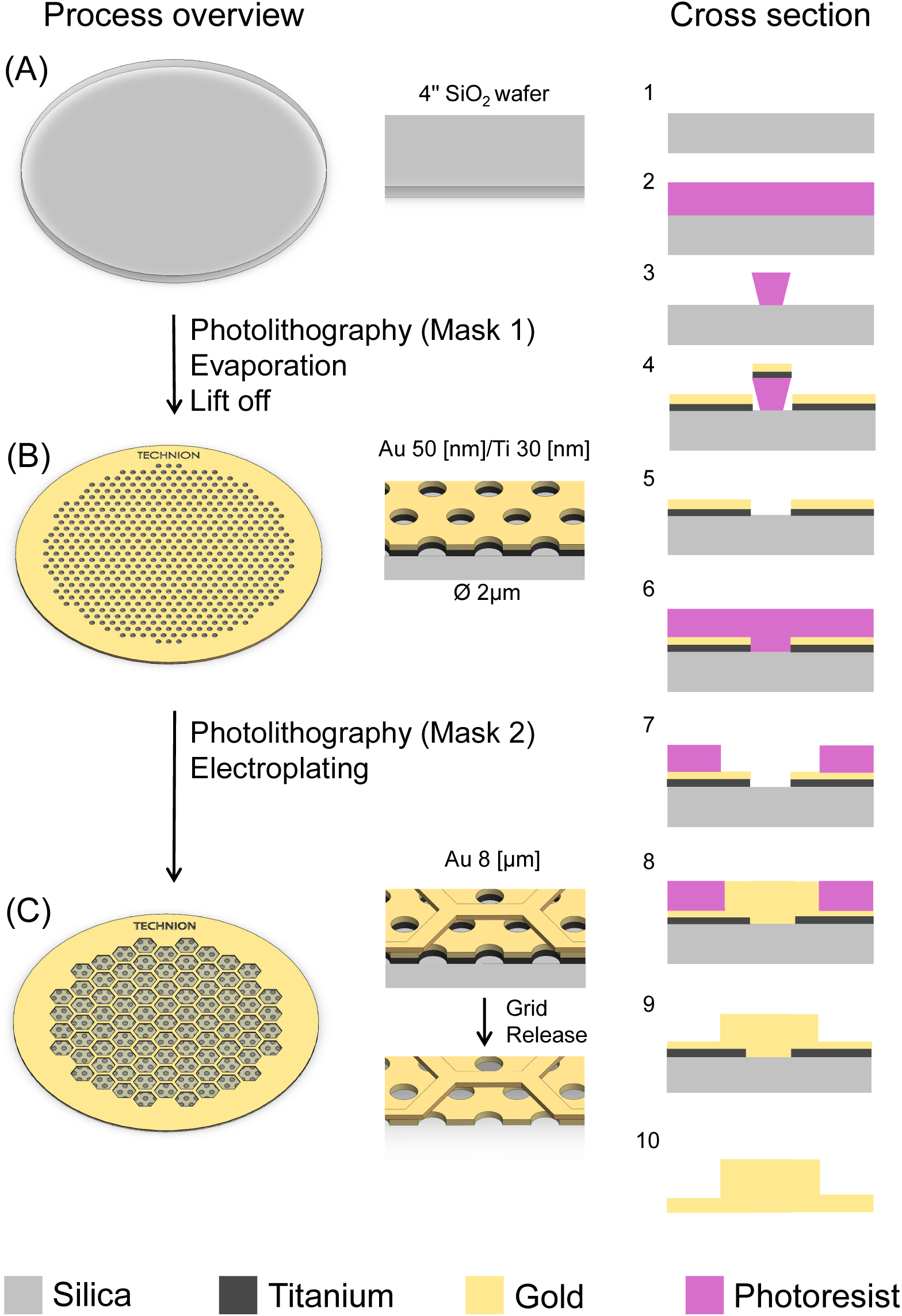
Wafer-scale process for manufacturing all-gold specimen supports for cryo-EM. (1) SiO_2_ wafer preparation. (2) Application of a positive resist operated in image reversal mode by spin coating. (3) Exposure of the wafer using a mask aligner to define the foil holes. (4) Metal evaporation to deposit a 30 nm titanium layer followed by a 50 nm gold layer. (5) Removal of the photoresist in NMP. (6) Spin coating of positive photoresist. (7) Multiple exposures through Mask 2 to define the grid mesh structure. (8) Electroplating 6.5–8 µm of gold on top of the seed layer. (9) Removal of the photoresist in NMP. (10) Wet etching of the titanium sacrificial layer and dissolving the SiO_2_ wafer in hydrofluoric acid to release the EM grids.

**Figure 2:**
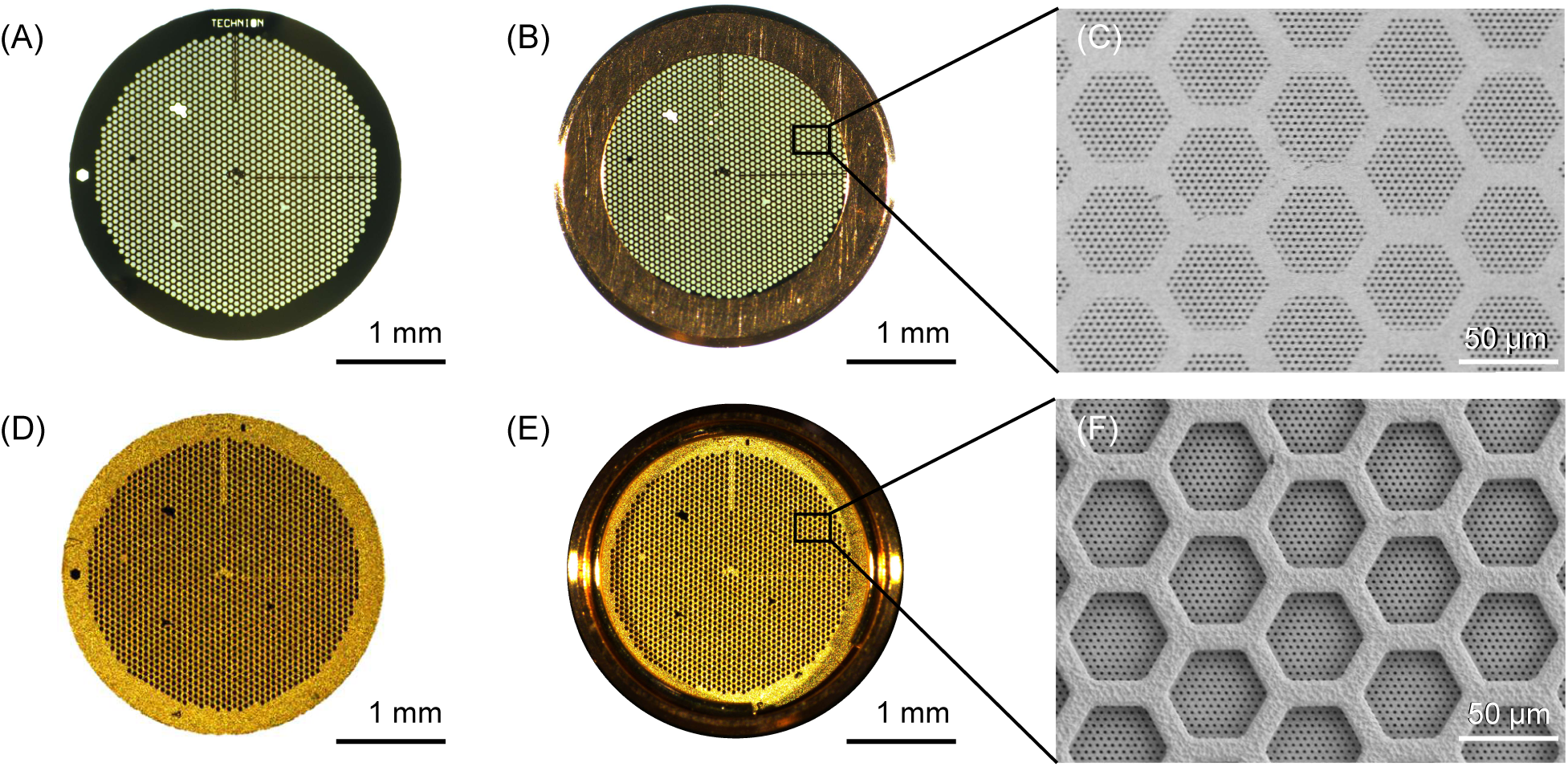
Batch-manufactured all-gold EM grids. (A) EM grid imaged from the foil side, with black areas representing the gold grid bars, gray areas representing the grid foil, and white areas indicating open regions. (D) Grid imaged from the back side. (B), (E) Grid clipped in a standard clip-ring/C-clip holder from the front (B) and back (E). (C), (F) Scanning electron micrographs of the grid from the foil side (C) and back side (F).

### Electroplating simulations tune grid-bar thickness and tilt access

To tune the electroplating process, we developed a numerical framework for modeling gold deposition during grid fabrication. The simulations capture the relationship between current density, deposition rate, and time-dependent gold growth as seed-layer holes fill during plating (Figures 3 and S1–S3). In practice, elapsed plating time was not a reliable predictor of final thickness because intermittent loss of electrical contact could occur during deposition. We therefore used the simulations to define an expected thickness window and to interpret geometry-dependent deposition trends, while the final grid-bar thickness was set and verified experimentally by stylus profilometry.

**Figure 3:**
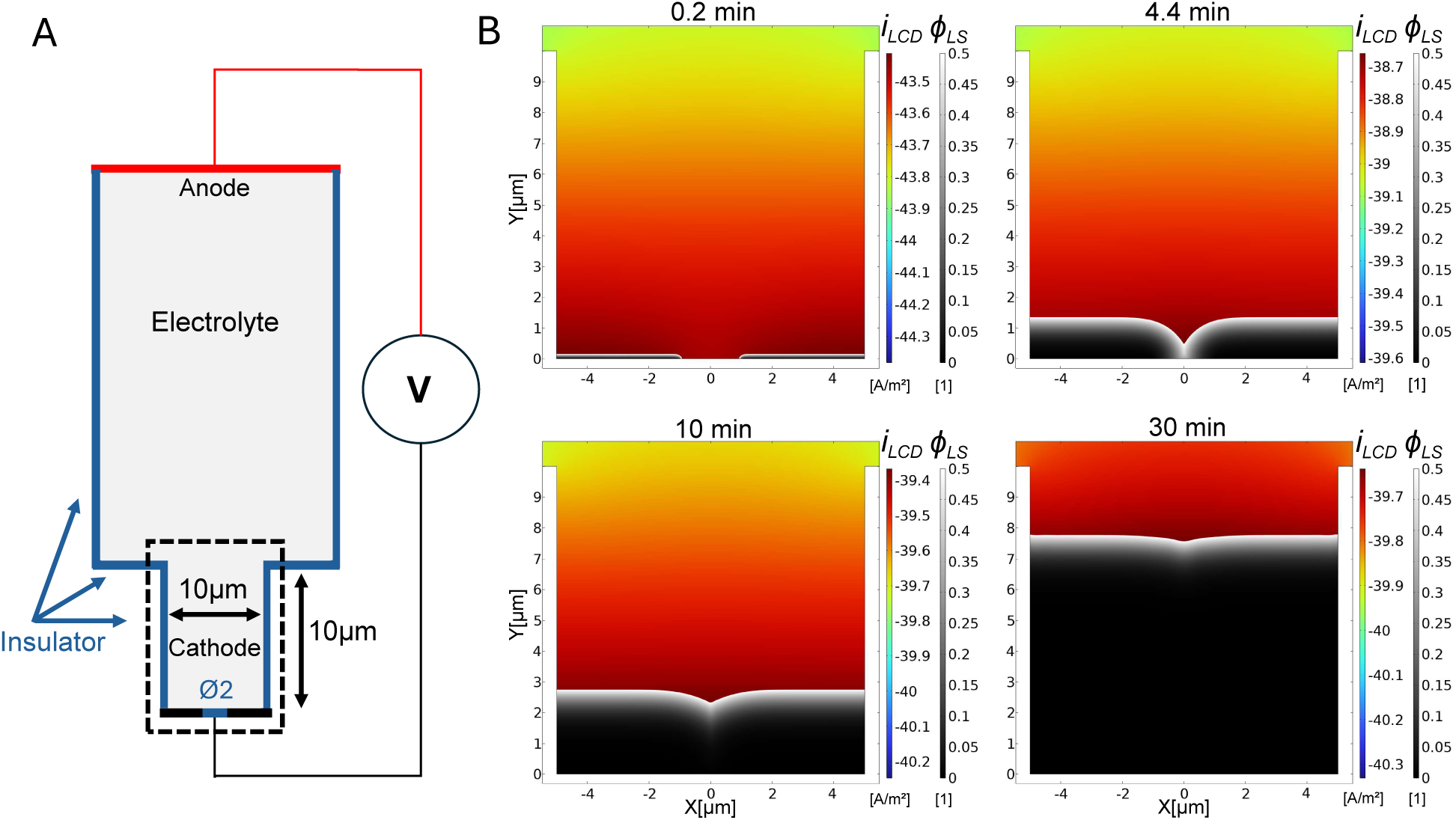
Two-dimensional numerical simulation of gold electroplating on a conductive surface around a 2 µm-diameter nonconductive hole. (A) Full computational domain used in the simulations, including the electrolyte reservoir, anode, cathode, and vertical symmetry boundaries. The trench region near the cathode, denoted by the dotted line, is shown in the simulation results. (B) Deposited gold (black) at 0.2, 4.4, 10, and 30 min. The local current density is denoted by *i*_LCD_, and the level-set variable by *ϕ*_LS_.

Profilometry showed structural bars in the 6.5–8 µm range (Figure S4), bracketing the 7.5 µm ideal predicted by the model and by the analytical Faraday estimate at the nominal deposition rate of 0.25 µm/min.^24^ This thickness window balances mechanical stability against electron transparency at high tilt. A 7–8 µm bar approximately permits tilt access of 73^◦^ (Figure S1), which is compatible with cryo-ET tilt-series acquisition. Although 2 µm foil holes weakly perturb the bulk deposition trajectory, the model captures the larger center-to-border thickness gap that develops for 6–8 µm holes (Figure S2), providing a quantitative framework for predicting and tuning gold thickness as a function of foil-hole geometry.

### Anisotropic foil geometries provide intrinsic contact-guidance cues

The versatility of the fabrication process enabled engineering of anisotropic foil geometries containing oval holes that provide intrinsic contact-guidance cues directly on the EM grid surface (Figures 4 and 5). The oval-hole and circular-hole grids were fabricated using the same wafer-scale process and shared the same mesh architecture, foil material, foil thickness, cell-culture conditions, staining protocol, and image-analysis pipeline; the programmed foil-hole geometry was the sole intended design variable. In contrast to chemical micropatterning, the anisotropic grids directed actin organization through substrate geometry alone. Cells cultured on grids containing oval holes exhibited preferential actin alignment along the major axis of the oval holes, whereas cells cultured on conventional circular-hole grids displayed a substantially more isotropic actin organization.

**Figure 4:**
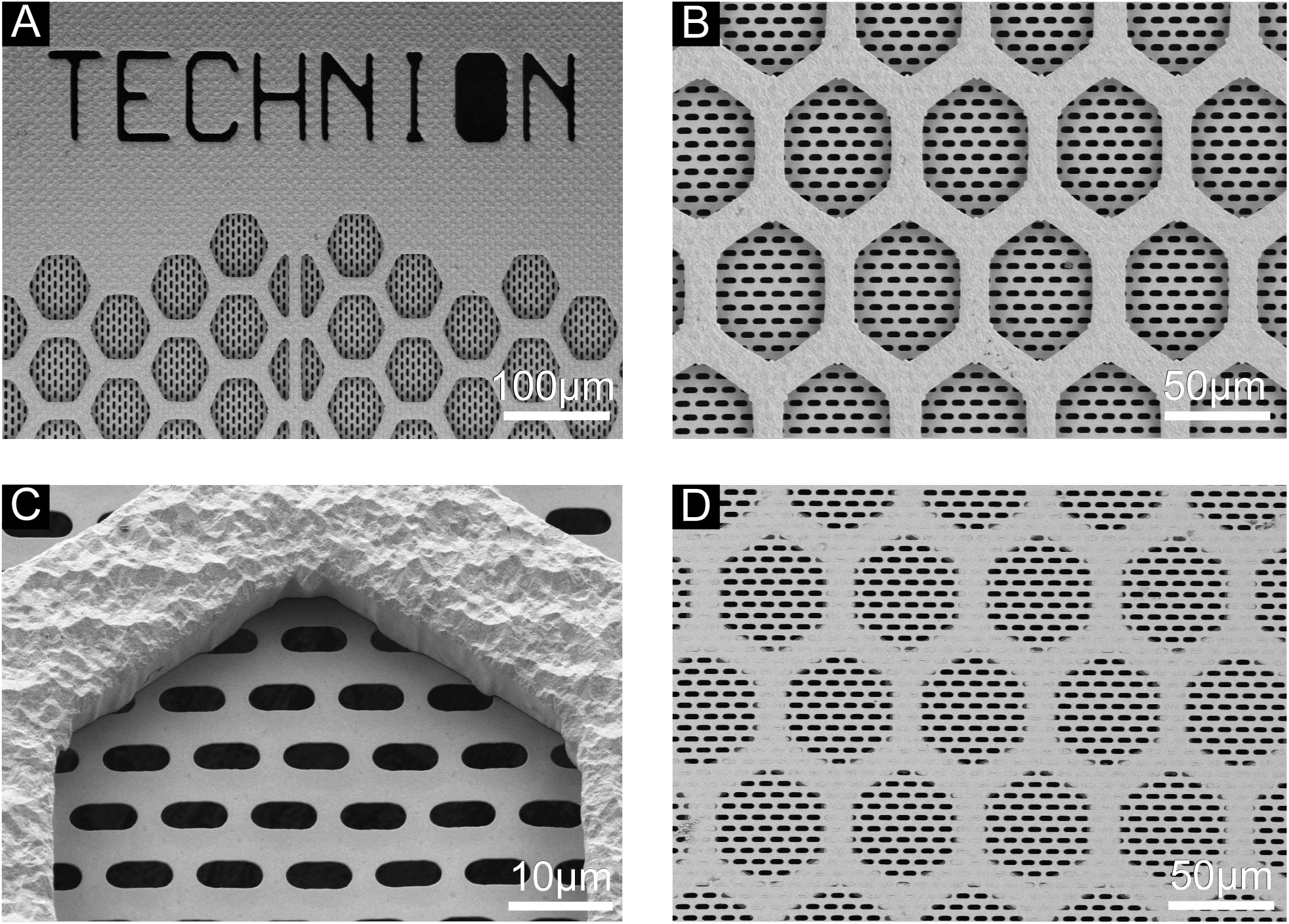
Scanning electron micrographs of batch-manufactured EM grids with oval gold foil holes. (A) Rim of the grid. (B) Back side of the grid. (C) Close-up of a single grid-bar hexagon imaged at a 35^◦^ tilt angle. (D) Front side of the grid.

**Figure 5:**
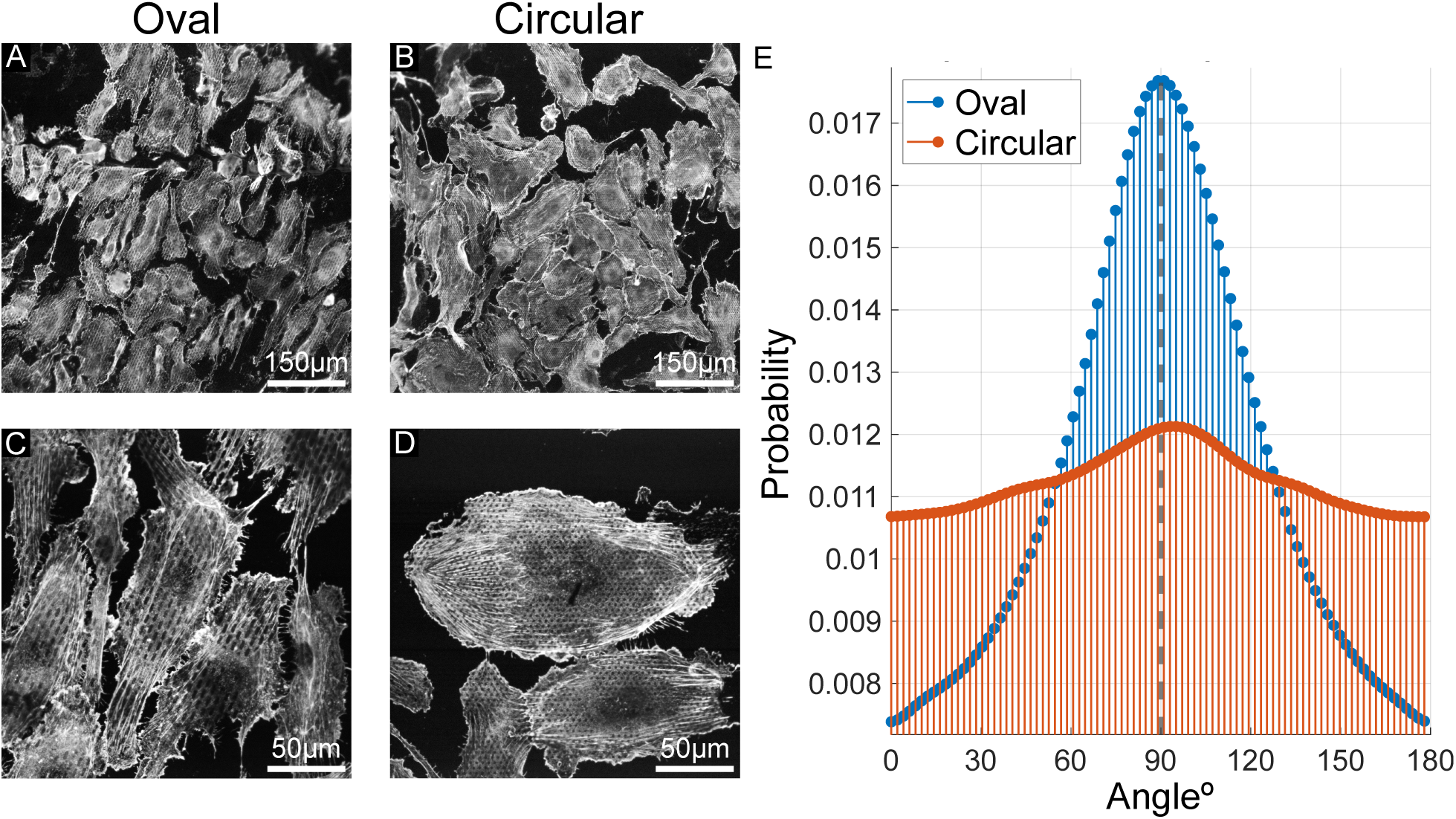
Actin orientation of cells cultured on anisotropic versus circular-hole grids. (A), (C) Cells cultured on EM grids with oval holes at 20× (A) and 60× (C) magnification. (B), (D) Cells cultured on EM grids with circular holes at 20× (B) and 60× (D) magnification. Actin is labeled with ActinRed 555. (E) Normalized orientation distributions of actin signal on EM grids. Curves show the normalized probability of local actin orientations for cells grown on oval and circular grids. Angles are referenced such that 90^◦^ (dashed line) is the oval major axis. The pronounced peak near 90^◦^ for oval grids indicates alignment along the major axis, whereas circular grids show a broader distribution without a distinct preferred angle. Grid-level AI statistics are reported in Table 1 and Table S1.

**Table 1:** Grid-level actin alignment index (AI) for oval-hole and circular-hole grids. Values are mean ± SD across grids (*n* = 4 grids per condition, sampled across 2 independent experiments). Welch’s *t*-test and Cliff’s *δ* were calculated on per-grid means. Cliff’s *δ* = 1.0 indicates complete rank separation: every oval-hole grid exceeded every circular-hole grid with no overlap. The exact Mann–Whitney *p*-value for this complete-separation ranking is 0.029.

| Half-window, $h$<br>(deg) | Oval AI<br>mean ± SD | Circular AI<br>mean ± SD | Welch $p$ | Cliff’s $\delta$ |
| --- | --- | --- | --- | --- |
| 10 | 0.1755 ± 0.0138 | 0.1206 ± 0.0055 | 0.0019 | 1.00 |
| 20 | 0.3328 ± 0.0234 | 0.2387 ± 0.0110 | 0.0015 | 1.00 |
| 30 | 0.4662 ± 0.0285 | 0.3533 ± 0.0150 | 0.0013 | 1.00 |

To quantify actin alignment, we computed an alignment index (AI) from local actin orientations measured using the Directionality plugin in ImageJ. Per-image AI values were averaged within each grid to yield one grid-level value, and group differences were evaluated using grid-level means from four oval-hole grids and four circular-hole grids (Table 1; Tables S1 and S2). The mean AI per grid was higher for oval grids than for circular grids at all angular window sizes, with complete rank separation between groups: every oval-hole grid had a higher mean AI than every circular-hole grid, with no overlap between conditions (Cliff’s *δ* = 1.0; Welch’s *t*-test *p <* 0.005; *n* = 4 grids per condition). The direction of the effect was consistent in both independent experiments, with oval-hole grids showing higher mean AI than circular-hole grids in each experiment separately. Analysis of the angular orientation distributions further supported this conclusion: the deviation magnitude from isotropy, computed from the pooled distributions, was approximately 7.2× greater for oval grids than for circular grids (Table S3). Together, the grid-level statistics and pooled angular distributions consistently demonstrate that anisotropic foil geometry strongly biases on-grid actin orientation.

These results demonstrate that EM support geometry alone can influence cellular organization. Anisotropic foil architectures, therefore, provide a simple physical strategy for guiding actin alignment that is integrated into the grid fabrication rather than added as a separate processing step, establishing the EM grid itself as an active, tunable biointerface rather than a passive support.

### Compatibility with cryo-EM and cellular sample-preparation workflows

To evaluate compatibility with high-resolution cryo-EM workflows, we performed SPA reconstruction of apoferritin using the fabricated all-gold grids. SPA imaging yielded a 2.15 Å global-resolution reconstruction on a 200 kV Glacios cryo-TEM, as estimated by the gold-standard Fourier shell correlation criterion at FSC = 0.143 (Figure 6). The reconstruction was obtained from 221,390 final particles selected from 971 micrographs, demonstrating that the grids support high-resolution cryo-EM imaging. Surface treatment with polyethylene glycol-thiol (PEG-SH) qualitatively improved grid wetting, allowing aqueous droplets to penetrate the foil holes more readily than on unmodified grids ^25^ (Figure S5).

**Figure 6:**
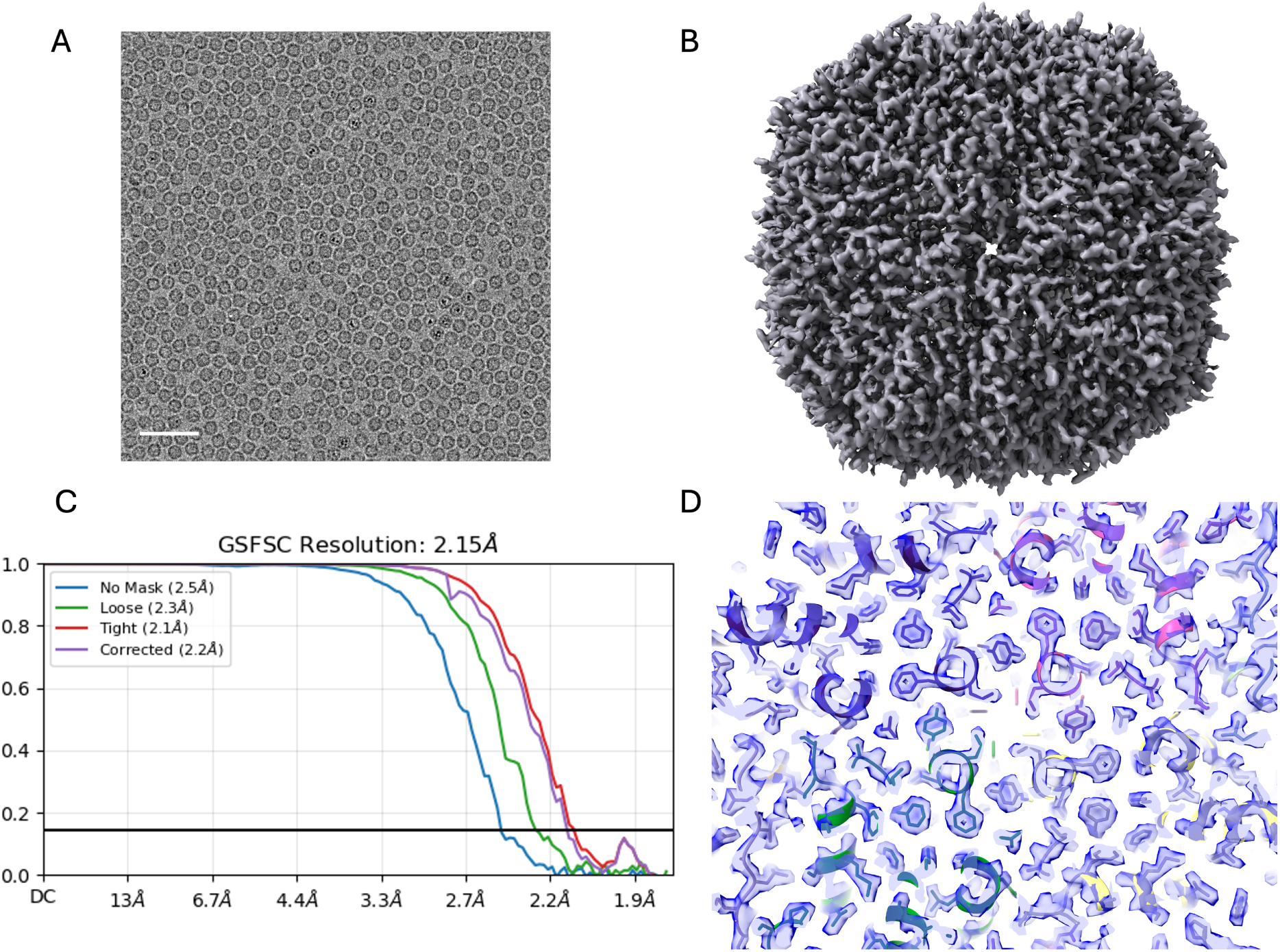
Single-particle analysis of apoferritin collected using all-gold EM grids. (A) Representative cryo-EM micrograph. Scale bar, 50 nm. (B) Cryo-EM map of apoferritin. (C) Gold-standard Fourier shell correlation (FSC) curve showing a global resolution of 2.15 Å at FSC = 0.143. (D) Close-up view of cryo-EM density (transparent blue) and fitted model at the fourfold symmetry axis.

We further evaluated compatibility with cellular sample-preparation workflows by performing protein micropatterning on the fabricated grids (Figure 7). The micropatterning process generated well-defined adhesive patterns that controlled cellular shape and positioning on the EM supports, a key preparatory step for cellular cryo-ET. The grids are also engineered for cryo-ET: the biocompatible all-gold structure, the 2 µm holes suited to imaging extended cellular regions, and the bar thickness optimized for high-tilt access (∼ 73^◦^, Figure S1) together satisfy the principal geometric requirements for tilt-series acquisition on vitrified cells.

**Figure 7:**
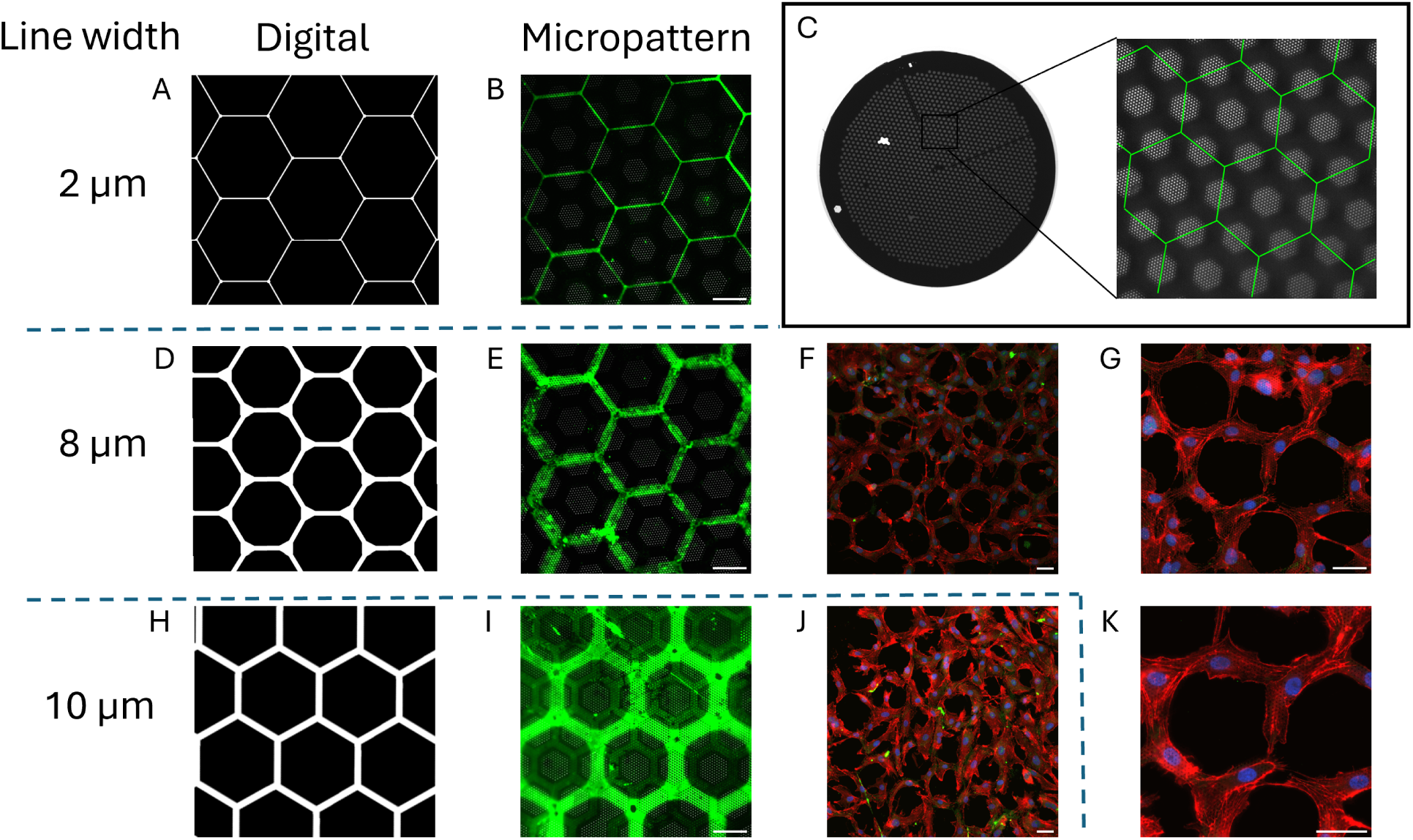
All-gold EM grids are compatible with micropatterning and cell culture. (A), (D), (H) Digital designs used to generate hexagonal micropatterns with line widths of 2, 8, and 10 µm. (B), (E), (I) Fluorescence micrographs of gelatin micropatterns on EM grids (green). (C) Schematic of a grid with a hexagonal micropattern (green). (F), (G), (J), (K) Fluorescence micrographs of cells cultured on the micropatterns, demonstrating successful cell attachment and organization. F-actin (red), nuclei (blue). All scale bars, 50 µm.

The present approach uses lift-off to define the foil geometry and titanium as a sacrificial layer, enabling grid release with a single etchant while maintaining an all-gold final support.

A limitation of the lift-off approach is that the minimum achievable hole diameter is constrained by the resolution of the photomask, which in our implementation was patterned by direct-write laser lithography (Heidelberg MLA-150) and limited to features of approximately 1 µm; wafer exposures were then performed by contact lithography (KARL SUSS MA-6). Although submicrometer holes are advantageous for minimizing beam-induced motion in SPA cryo-EM,^9^ larger foil holes are well suited for cryo-ET applications, where collecting a tilt series over larger cellular regions is often desirable.

## Conclusions

We developed a wafer-scale, all-standard-cleanroom process for fabricating customizable all-gold EM grids without individual grid handling. The process produces 594 grids per 4-inch wafer, supports high-resolution SPA cryo-EM at 2.15 Å resolution, and is compatible with cellular micropatterning workflows. Its central advantage is architectural flexibility: by changing the foil-hole design, the grid can be converted from a passive specimen support into an active physical cue that biases cellular organization without requiring any additional surface-chemistry steps. Oval-hole grids induced robust actin alignment relative to circular-hole controls, with complete grid-level separation across all alignment-index windows tested. These results establish anisotropic all-gold grids as a practical platform for coupling cryo-EM specimen support, microfabrication scalability, and geometry-based control of cellular organization. Future work should extend this approach to additional foil geometries, cell types, and in-cell tomography workflows.

## Methods

### Overview of grid fabrication

The process we developed for fabricating all-gold EM grids requires two lithography steps, metal deposition by evaporation and electroplating, and wet etching. The fabrication steps are summarized in Figure 1 and described below.

### Fabricating the holey foil

The process began with a fused-silica wafer that was micropatterned with a thin Ti/Au film by lift-off to define the grid foil (Figure 1A,B). The chrome photomasks used here (Mask 1A for round 2 µm holes and Mask 1B for oval 2 µm × 6 µm holes; Figures S6 and S7) were written by direct-write laser lithography (Heidelberg MLA-150). The wafer was coated with a positive photoresist operated in image reversal mode (AZ5214) by spin coating at 5000 rpm for 60 s, soft-baked at 110 ^◦^C for 1.5 min, and exposed through the photomask in a contact mask aligner (KARL SUSS MA-6). A post-exposure bake was performed at 120 ^◦^C for 60 s, followed by a 10 s flood exposure and development, generating an undercut resist profile for lift-off. The wafer was then metallized by evaporation (Evatec BAK-501A) with a 30 nm titanium adhesion/sacrificial layer followed by a 50 nm gold seed layer. After metal deposition, the photoresist was dissolved in *N* -methyl-2-pyrrolidone (NMP) overnight, and residual lifted-off metal flakes were removed by rinsing with acetone.

### Defining the grid bars

The grid perimeter and grid bars were defined lithographically using thick positive photoresist. AZ4562 was spin-coated at 1250 rpm for 60 s and baked at 110 ^◦^C for 7 min. The wafer was exposed in the mask aligner using multiple exposures to improve degassing, followed by a postexposure bake at 90 ^◦^C for 2 min. Allowing sufficient time between exposure and hard bake improved resist quality, consistent with outgassing of nitrogen from the photoresist layer before baking. To ensure mechanical stability of the grids, the photoresist layer must be sufficiently thick to support the targeted electroplated gold thickness; profilometry confirmed a resist height of approximately 12.5 µm, exceeding the targeted plated thickness (Figure S4). Alternative routes to a thick resist coating include double coating or the use of a thicker photoresist compatible with electroplating.

### Electroplating

A thick gold layer was deposited into the trenches defined by the photoresist, using the conductive thin gold foil as the seed layer. The patterned wafer served as the cathode, while a platinum electrode served as the anode. We used a commercially available gold sulfite/thiosulfite electrolyte solution (MetGold ECF 33B) and electroplating parameters in the optimal range recommended by the manufacturer: a current density of 4 mA/cm^2^ at 54 ^◦^C, corresponding to a nominal deposition rate of approximately 0.25 µm/min.

### Grid release

To release the grids from the wafer, the photoresist was first removed in NMP. The fused silica and titanium layers were then dissolved in 49% hydrofluoric acid (HF). *Caution:* HF is highly toxic and corrosive; all HF handling was performed according to institutional safety procedures using compatible protective equipment, appropriate fume-hood containment, and calcium gluconate gel availability. The thin foil connectors between grids broke spontaneously during release, leaving individual grids in solution. A custom Teflon holder with holes around the base was designed to allow immersion of the entire wafer in HF while retaining released grids and enabling rinsing in water (Figure S8). Released grids were stored either in water or dried and stored in grid boxes. Note: residual gold flakes generated during lift-off can contaminate the HF bath.

### Grid sorting and storage

Drying released grids is complicated by capillary aggregation as water drains. Individual grids can be removed manually and placed on filter paper for drying, but this procedure is labor-intensive. To improve throughput, we designed a custom 3D-printed grid-sorting device that uses radial symmetry to separate grids during drying (Figure S9). A porous base contains holes small enough to retain the grids while allowing water to drain. We also designed grid boxes with space for 4–12 grids that fit into 15 or 50 mL tubes for grid functionalization and storage (CAD files are provided in the Supporting Information).

### Functionalization for SPA

To improve wetting before SPA sample application, grids were coated with polyethylene glycol-thiol (PEG-SH). After 1 min exposure to atmospheric plasma, the grids were incubated in 1 mg/mL PEG-SH (Sigma-Aldrich, 729140) in ultrapure water for 1 h. Wettability was assessed qualitatively by placing a 7 µL droplet of deionized water on a vertically held grid with fine-point tweezers and comparing droplet penetration through functionalized and unmodified foils^25^ (Figure S5).

### Micropatterning

For micropatterning, we followed the process previously used on commercially available grids.^18–20^ Briefly, grids were exposed to atmospheric plasma for 1 min and incubated overnight in 0.01% poly-L-lysine (Sigma-Aldrich, P4707), followed by a 1 h incubation in PEG-SVA (Laysan Bio) to generate an antifouling coating. PEG was then selectively degraded using the Alveole PRIMO maskless UV exposure system with PLPP gel (Nanoscale Labs) as the photoinitiator.

### Cell culture and immunofluorescence

Human umbilical vein endothelial cells (HUVECs; ScienCell, 8000-SCL) were cultured in EGM-2 MV Microvascular Endothelial Cell Growth Medium supplemented with penicillin, streptomycin, and growth factors. Cells were maintained at 37 ^◦^C and 5% CO_2_.

Before seeding, micropatterned grids were back-filled with 0.2% gelatin–Oregon Green 488 conjugate (Thermo Fisher Scientific, G13186) for 1 h at room temperature. Unpatterned grids were exposed to atmospheric plasma for 1 min and incubated with 0.2% gelatin solution (Sigma-Aldrich, G1393). To seed cells on the grids, 5 µL of a cell suspension of approximately × 10^6^ cells/mL was added to each grid. After cells began adhering to the grids (∼1 h), mL of warm medium was added to each dish, and cells were incubated overnight at 37 ^◦^C and 5% CO_2_. Cells were fixed in 4% paraformaldehyde (PFA; Rhenium, 043368.9M) in PBS.

For immunostaining, cells were permeabilized using antibody dilution buffer (ADB). Following a 1 h blocking step with ADB, cells were stained with ActinRed 555 (Thermo Fisher Scientific, R37112) and Hoechst 33342. Cells were imaged on a Nikon ECLIPSE Ti2-E spinning-disk microscope using 20× air, 40× oil, and 60× oil-immersion objectives.

### Statistical analysis of actin orientation

To assess whether grids with oval holes bias actin orientation, cells were cultured on oval-hole grids and standard circular-hole grids, and actin orientation was quantified from fluorescence images. Orientation distributions were computed with the Directionality plugin in ImageJ/Fiji using the local-gradient orientation method.^26,27^ Because orientation is axial, angles were reported on [0, 180). All images were referenced so that 90^◦^ denotes the oval major axis and 0^◦^*/*180^◦^ denotes the perpendicular direction (Figure 5). Histograms were computed with ∼2^◦^ bins and, for display only, circularly smoothed with an area-normalized Gaussian kernel (*σ* = 6^◦^) to reduce high-frequency noise.

Anisotropy of actin orientations was quantified as the mean squared deviation of the angular probability distribution from a uniform distribution.^28^ It is defined as

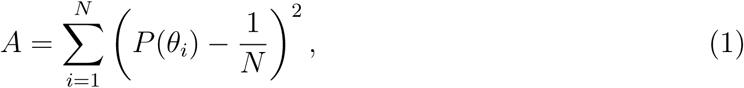

where *P* (*θ_i_*) is the normalized probability of observing an actin orientation in the *i*th angular bin centered at *θ_i_*, with

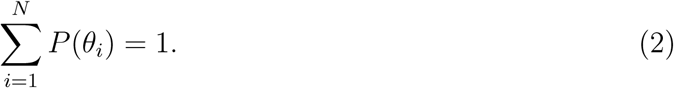

Here, *θ_i_* denotes the actin orientation angle measured with respect to the reference direction in the *X*-*Y* plane. A value of *A* = 0 corresponds to a perfectly isotropic angular distribution, while larger values indicate increasing directional anisotropy.

An alignment index (AI) was computed for each image histogram to quantify how strongly the local orientation aligns with the substrate direction. Let *w*(*θ*) denote the gradient-weighted orientation counts at angle *θ* in degrees. For a half-window *h* around the substrate axis (90^◦^),

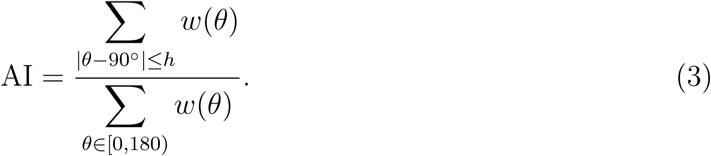

AI ranges from 0, indicating little signal near 90^◦^, to 1, indicating that all the signal is concentrated near 90^◦^. A higher AI indicates stronger alignment of actin orientation with the oval major axis. Per-image AI values were averaged within each grid to yield one grid-level value. Group differences were evaluated using Welch’s two-sample *t*-test on grid-level means from four oval-hole grids and four circular-hole grids collected across two independent experiments. Effect size was quantified using Cliff’s *δ* on the per-grid means (Table S1).

### Gold electroplating simulation

Precise control of the electroplated layer thickness is important for grid performance. During electron tomography, the grid is tilted over a wide angular range to acquire projections for 3D reconstruction. If the supporting gold layer is excessively thick, it can obstruct the electron beam at high tilt angles, whereas a layer that is too thin compromises mechanical stability. This trade-off between robustness and transparency requires control of deposited thickness.

In our fabrication process, the seed layer is patterned with 2 µm holes that fill with gold over time, producing nonlinear deposition dynamics. To predict and interpret the electroplating process in this geometry, we developed COMSOL Multiphysics models of gold electroplating (Figure S1). The simulations used phase-field and level-set methods to capture the time-dependent evolution of gold deposition (Figures S2 and S3). Simulation parameters are reported in Table S4. As deposition proceeds, the effective plating area increases as foil holes fill, leading to a corresponding reduction in current density.^29^

In the model, gold is deposited onto the cathode surface, which evolves dynamically as deposition progresses. We assume a well-stirred electrolyte, constant ionic concentration outside the diffusion layer, and mass transport at the cathode driven by diffusion and electromigration.

### Cryo-EM sample preparation

Three microliter aliquots of apoferritin in solution were deposited on glow-discharged 2 µm-hole grids before manual blotting for 2 s at room temperature. The grids were vitrified by rapid plunging into liquid ethane using a home-built plunging apparatus. Frozen samples were stored in liquid nitrogen until imaging.

### Cryo-EM data acquisition

Cryo-EM data were collected at cryogenic temperature on a Glacios microscope equipped with a Falcon 4i direct electron detector and a Selectris X energy filter (Thermo Fisher Scientific). The microscope was operated at 200 kV, and the energy filter was set to ±5 eV from the zero-loss peak. Movies were recorded in dose-fractionated counting mode using EPU (Thermo Fisher Scientific) with a pixel size of 0.89 Å, a total electron dose of 30 *e*^−^*Å*^2^ a defocus range of −0.5 to −1.5 µm. Data collection parameters are reported in Table S5.

### Cryo-EM data processing

Dose-fractionated image stacks (1255 total) were imported into cryoSPARC v4.7.0.^30^ Movies were subjected to patch motion correction and patch contrast transfer function (CTF) estimation. A cutoff of CTF fit resolution better than 5.0 Å was applied, resulting in 971 micrographs used for further processing. Template picker was used for particle picking. A total of 506,107 particles were extracted with a box size of 300 pixels and subjected to one round of two-dimensional classification, yielding 221,390 particles selected for further refinement. The final map had an average resolution of 2.15 Å, estimated using the gold-standard Fourier shell correlation criterion at FSC = 0.143.

## Supporting Information Available

Supporting Information: Supplementary methods, photomask layouts, profilometry data, PEG-SH wetting assessment, cryo-EM acquisition and processing parameters, and actin-orientation statistics, CAD files for the custom release, sorting, and storage devices.

## Data Availability Statement

Data are provided as Supporting Information or Review-Only Material. Additional raw data are available from the corresponding author upon reasonable request.

## Author Contributions

A.A. and L.E. conceived the study and designed the fabrication process. A.A. performed microfabrication. N.B.A. performed cell culture and fluorescence imaging experiments. R.Z. performed cryo-EM data acquisition and data processing. A.A. performed electroplating simulations and statistical analysis of actin orientation. L.E. supervised the project. All authors contributed to writing the manuscript and approved the final version.

## Acknowledgement

This study was inspired by the pioneering work of Chris Russo (LMB). Microfabrication was performed at the Technion Micro-Nano Fabrication Unit (MNFU), and the authors thank MNFU staff members A. Shacham and Y. Milyutin for their assistance. This work was partially supported by Israel Science Foundation (ISF) Personal Research Grant no. 1925/23 and by a grant from the Diane and Guilford Glazer Foundation (L.E.). We thank O. Kleinerman for HRSEM imaging, which was performed at the Technion Center for Electron Microscopy of Soft Matter. We thank G. Kozyukin from the Israel Institute of Materials Manufacturing Technologies for electroplating advice. The authors are grateful for the generous support from the Guzik Foundation to Ben-Gurion University’s cryo-electron microscopy unit, where cryo-EM was performed. AI was used for language-editing assistance, formatting assistance, and preparation. The authors reviewed and edited all content and take full responsibility for the manuscript.

## TOC Graphic

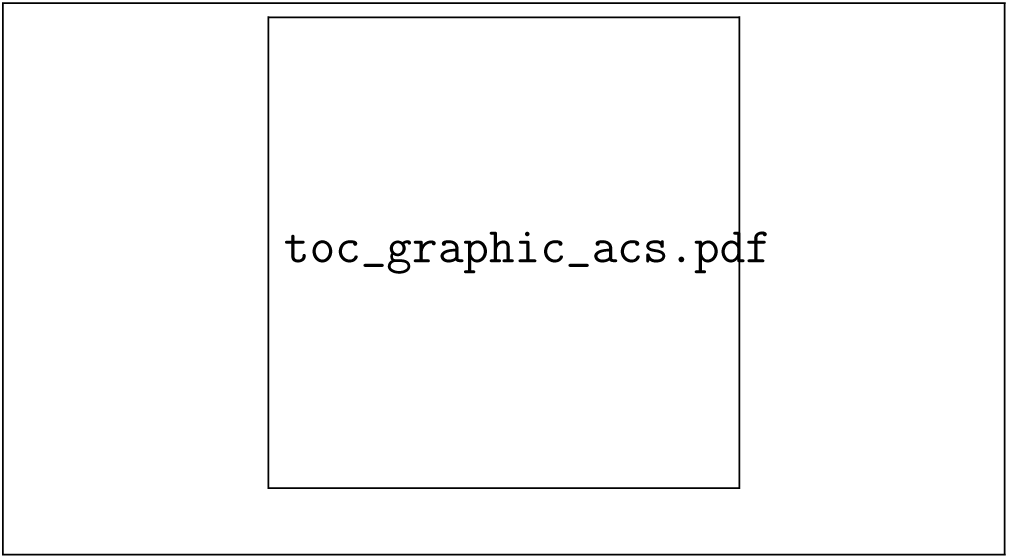

