## Supplemental Information for "Gold Electron Microscopy Grids with Anisotropic Foil Geometry Enable On-Grid Contact Guidance"

for

**S2 Supplementary Figures** **5**

**S3 Supplementary Tables** **14**

**S4 Supporting Equations** **19**

### S1 Supplementary Methods

#### S1.1 Materials

Fabrication of the all-gold grids used 4-inch fused-silica wafers, AZ5214 negative/image-reversal photoresist for foil patterning, AZ4562 positive photoresist for the electroplating pattern, titanium and gold for evaporation, N-methyl-2-pyrrolidone (NMP) for lift-off and resist stripping, MetGold ECF 33B gold sulfite/thiosulfite electrolyte for electroplating, a platinum anode, and hydrofluoric acid (HF, 49%) for release of the finished grids.

Surface modification and biological experiments used PEG-thiol for single-particle analysis (SPA) functionalization (PEG-SH, Sigma-Aldrich, 729140), poly-L-lysine (Sigma-Aldrich, P4707), PEG-SVA (Laysan Bio), PLPP gel photoinitiator (Nanoscale Labs), gelatin (Sigma-Aldrich, G1393), gelatin–Oregon Green 488 conjugate (Thermo Fisher Scientific, G13186), ActinRed 555 (Thermo Fisher Scientific, R37112), Hoechst 33342, paraformaldehyde (Rhenium, 043368.9M), and HUVECs (ScienCell, 8000-SCL).

#### S1.2 Grid fabrication, electroplating, release, and handling

All-gold grids were fabricated on 4-inch fused-silica wafers using a two-mask process. For the holey foil, the wafer was coated with AZ5214 at 5000 rpm for 60 s and soft-baked at 110 °C for 1.5 min. After exposure through the foil mask, the wafer underwent a postexposure bake at 120 °C for 60 s followed by a 10 s flood exposure before development. The patterned wafer was metallized by evaporation of a 30 nm Ti adhesion/sacrificial layer and a 50 nm Au seed layer, and unwanted metal was removed by overnight NMP lift-off. Depending on the mask, the foil contained either 2 µm circular holes or anisotropic oval holes measuring 2 µm by 6 µm.

The structural mesh was defined in AZ4562 spun at 1250 rpm for 60 s and baked at 110 °C for 7 min. Exposure was carried out in multiple passes to reduce degassing artifacts, and the patterned wafer was post-baked at 90 °C for 2 min. Stylus profilometry confirmed that the photoresist pattern height was approximately 12.5 µm, exceeding the targeted plated thickness. Gold bars were electroplated from the thin Au foil seed layer using the cyanide-free MetGold ECF 33B sulfite/thiosulfite electrolyte with a platinum counter electrode. The nominal plating conditions were 4 mA $cm^{-2}$ and 54 °C, corresponding to an analytical deposition rate of approximately 0.25 µm $min^{-1}$. Because nucleation is not instantaneous and intermittent electrical disconnection can occur during plating, elapsed plating time was not treated as the primary control variable. Instead, bar thickness was verified by profilometry after fabrication; representative grids showed structural bars in the 6.5–8 µm range, with slightly higher deposition toward the wafer edge.

After electroplating, the photoresist was stripped in NMP, and the fused-silica wafer together with the Ti sacrificial layer was dissolved in 49% HF to release individual grids. A custom Teflon holder was used to support the wafer during immersion and transfer between HF and deionized water. Released grids were transferred to deionized water, separated with a radial-flow sorting device to prevent aggregation during drying, and either dried in air or kept in water until use. Grid boxes accommodating 4–12 grids were also designed to fit 15 and 50 mL tubes for functionalization and storage.

#### S1.3 Surface functionalization, wetting assay, and micropatterning

For SPA applications, grids were exposed to atmospheric plasma for 1 min and then incubated in 1 mg $mL^{-1}$ PEG-SH in ultrapure water for 1 h to form a thiol-based self-assembled monolayer on the gold surface. Wettability was assessed qualitatively by placing a 7 µL droplet of deionized water on a vertically held grid with fine-point tweezers and comparing droplet penetration through functionalized and unmodified foils.

Micropatterning was performed using a previously established EM-grid workflow. Plasma-treated grids were incubated overnight in 0.01% poly-L-lysine and then for 1 h in PEG-SVA to create an antifouling coating. Hexagonal adhesive patterns were generated by maskless UV exposure on a PRIMO system using PLPP gel as the photoinitiator. Pattern fidelity and cell guidance on representative hexagonal designs are shown in the main manuscript; the supplementary figures focus on fabrication, validation, wetting, simulation, and orientation-analysis details not shown in the main text.

#### S1.4 Cell culture, fixation, and fluorescence imaging

HUVECs were cultured in EGM-2 MV Microvascular Endothelial Cell Growth Medium supplemented with penicillin, streptomycin, and growth factors. Cells were maintained at 37 °C and 5% $CO_2$ on dishes precoated with 0.2% gelatin.

Before seeding, micropatterned grids were back-filled for 1 h at room temperature with 0.2% gelatin–Oregon Green 488 conjugate. Unpatterned grids were plasma-treated for 1 min and incubated with 0.2% gelatin solution. To seed cells on the grids, 5 µL of a suspension containing approximately $10^6$ cells $mL^{-1}$ was added to each grid. After approximately 1 h of initial adhesion, 2 mL of warm medium was added, and the grids were incubated overnight. Cells were fixed in 4% paraformaldehyde in PBS, permeabilized with antibody dilution buffer, blocked for 1 h, and stained with ActinRed 555 and Hoechst 33342. Fluorescence imaging was performed on a Nikon ECLIPSE Ti2-E spinning-disk microscope using 20× air, 40× oil, and 60× oil-immersion objectives.

#### S1.5 Orientation analysis and statistics

To test whether anisotropic oval-hole grids bias cytoskeletal organization, actin images acquired from cells on oval-hole and circular-hole grids were analyzed in Fiji/ImageJ using the Directionality plugin with the local-gradient-orientation method. Because actin-orientation measurements are axial, angles were represented on the interval $[0, 180)$ rather than $[0, 360)$. All images were referenced so that 90° corresponded to the long axis of the oval holes and 0°/180° corresponded to the orthogonal direction. Histograms were computed with approximately 2° bins and, for display only, circularly smoothed with an area-normalized Gaussian kernel with $\sigma = 6°$.

Two independent experiments were analyzed. Each experiment included two oval-hole and two circular-hole grids, yielding four grids per condition in total. Eight to eleven whole-field images were acquired per oval-hole grid and seven to ten whole-field images were acquired per circular-hole grid, for a total of 38 oval-hole images and 36 circular-hole images. Alignment indices (AI) were computed on a per-image basis for half-windows $h = 10°$, 20°, and 30° about the substrate axis. For statistical testing, per-image AI values were first averaged within each grid, yielding one grid-level mean per grid ($n = 4$ grids per condition). Group differences were evaluated on these grid-level means with Welch's two-sample $t$-test, and effect size was reported using Cliff's $\delta$.

#### S1.6 Electroplating model and numerical implementation

Two-dimensional finite-element simulations were performed in COMSOL Multiphysics 6.3 using the Electrochemistry Module. The model combined the Tertiary Current Distribution, Nernst–Planck formulation with either phase-field (PF) or level-set (LS) interface tracking to simulate gold electrodeposition in a trench geometry approximating the experimental seed layer. The full numerical domain comprised an electrolyte reservoir with the anode at the upper boundary, the cathode at the lower boundary, and insulating vertical boundaries. For clarity, the result figures show only the trench region near the cathode.

Electrolyte transport was assumed to occur by diffusion and electromigration in a well-stirred sulfite-based electrolyte with constant ionic concentration outside the diffusion layer. The modeled reaction sequence was

$$[\mathrm{Au(SO_3)_2}]^{3-} \rightarrow [\mathrm{Au(SO_3)}]^{-} + \mathrm{SO_3^{2-}}, \tag{S1}$$

$$[\mathrm{Au(SO_3)}]^{-} \rightarrow \mathrm{Au^{+}} + \mathrm{SO_3^{2-}}, \tag{S2}$$

$$\mathrm{Au^{+}} + \mathrm{e^{-}} \rightarrow \mathrm{Au}. \tag{S3}$$

The interface-thickness parameter was set to $\varepsilon = 335$ nm in both PF and LS formulations. Quartic triangular elements and adaptive mesh refinement near the moving interface were used to ensure numerical stability and mesh-independent predictions. Mesh convergence was verified on a 2 μm hole geometry. For reporting pinch-off and convergence, the deposited region was identified using $\phi_{\mathrm{PF}} \geq 0.9$ in the phase-field formulation and $\phi_{\mathrm{LS}} \leq 0.1$ in the level-set formulation.

#### S1.7 Cryo-EM specimen preparation, acquisition, and processing

Apoferritin was used as a benchmark specimen for cryo-EM compatibility.

Protein solution was applied to grids with 2 μm circular holes, manually blotted for 2 s at room temperature to form a thin liquid film spanning the holes, and plunge-frozen in liquid ethane using a home-built plunging apparatus. Frozen grids were stored in liquid nitrogen until imaging.

Cryo-EM data were collected on a Thermo Fisher Glacios 200 kV cryo-TEM equipped with a Falcon 4i direct electron detector and a Selectris X energy filter. The energy-filter slit was set to ±5 eV from the zero-loss peak. Movies were acquired in dose-fractionated counting mode using EPU with a pixel size of 0.89 Å, a total exposure of 30 $e^{-}$ Å$^{-2}$, and a defocus range of $-0.5$ to $-1.5$ μm. A total of 1255 movies were imported into cryoSPARC v4.7.0 and processed by patch motion correction and patch CTF estimation. After applying a CTF fit-resolution cutoff better than 5.0 Å, 971 micrographs were retained. Template picking yielded 576,660 extracted particles, and one round of two-dimensional classification yielded 221,390 particles for refinement. The final reconstruction reached a global resolution of 2.15 Å by gold-standard Fourier shell correlation at FSC = 0.143.

For cryo-electron tomography trials, endothelial cells were cultured overnight on anisotropic grids, plunge-frozen on a Leica GP2 with a 13 s blot time, clipped, and loaded into the Glacios platform for tilt-series acquisition. Ice thickness was not suitable for interpretable tomography under the conditions tested; tomographic data are therefore not reported as a result in this manuscript.

### S2 Supplementary Figures

Figures S1–S9 provide the analytical framework, fabrication validation, wetting, orientation-analysis details, and custom devices that support the main text while avoiding duplication of the primary manuscript figures.

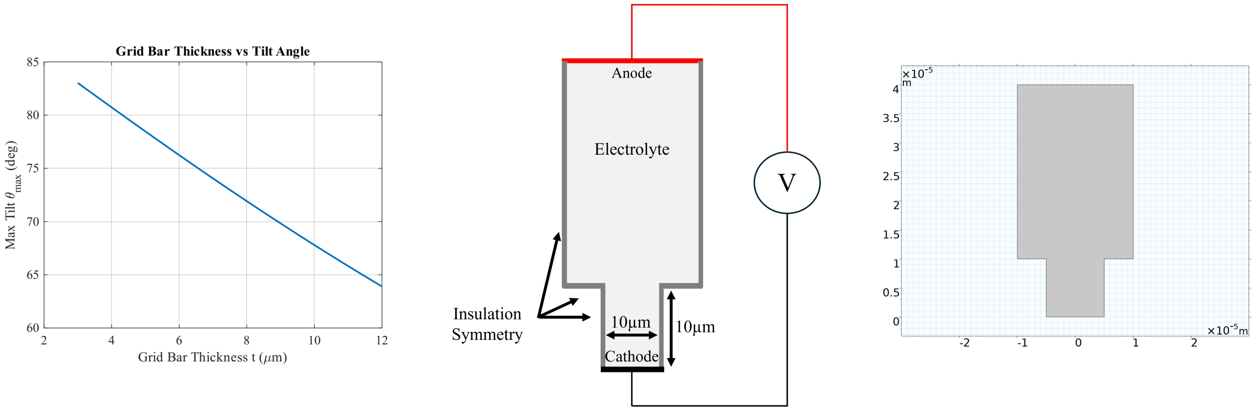

(A) (B) (C)

Figure S1: Analytical and numerical framework for electroplating simulations. (A) Maximum accessible tilt angle as a function of grid-bar thickness; a 7–8 μm structural layer corresponds to approximately 73° tilt access. (B) Full electrochemical reservoir used in the COMSOL model. (C) Computational geometry as depicted in the COMSOL interface.

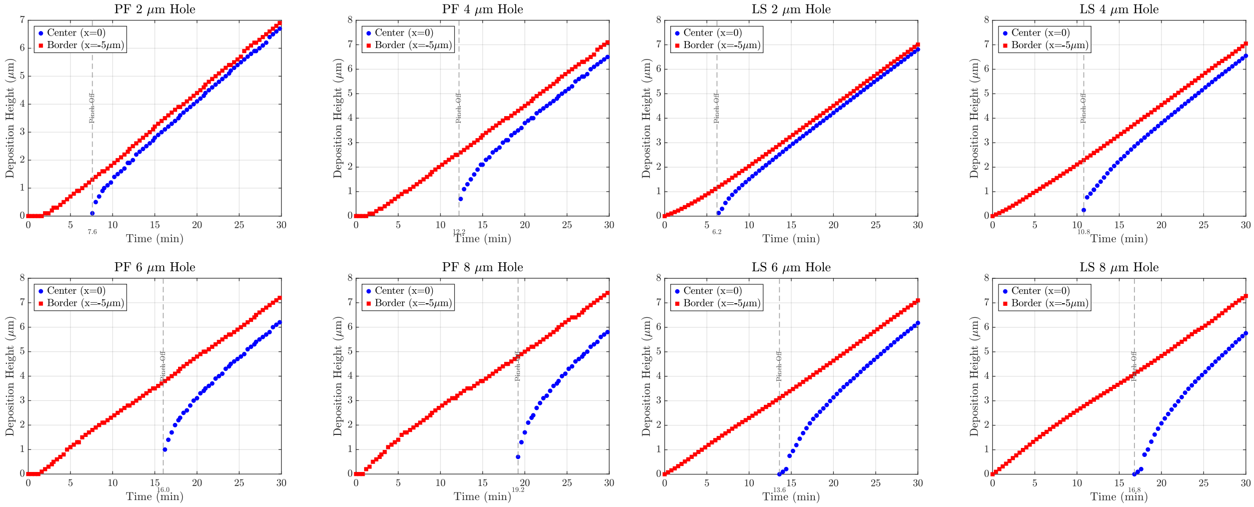

(A) (B)

Figure S2: Comparison of phase-field (PF) and level-set (LS) predictions for hole filling across seed-layer hole diameters. (A) PF formulation. (B) LS formulation. Red points denote deposition thickness at the trench border, and blue points denote thickness at the hole center; vertical dashed lines mark pinch-off. In both formulations, 2 and 4 µm holes rapidly approach the bulk deposition trajectory after closure, whereas 6 and 8 µm holes retain a larger center-to-border gap over the simulated time window.

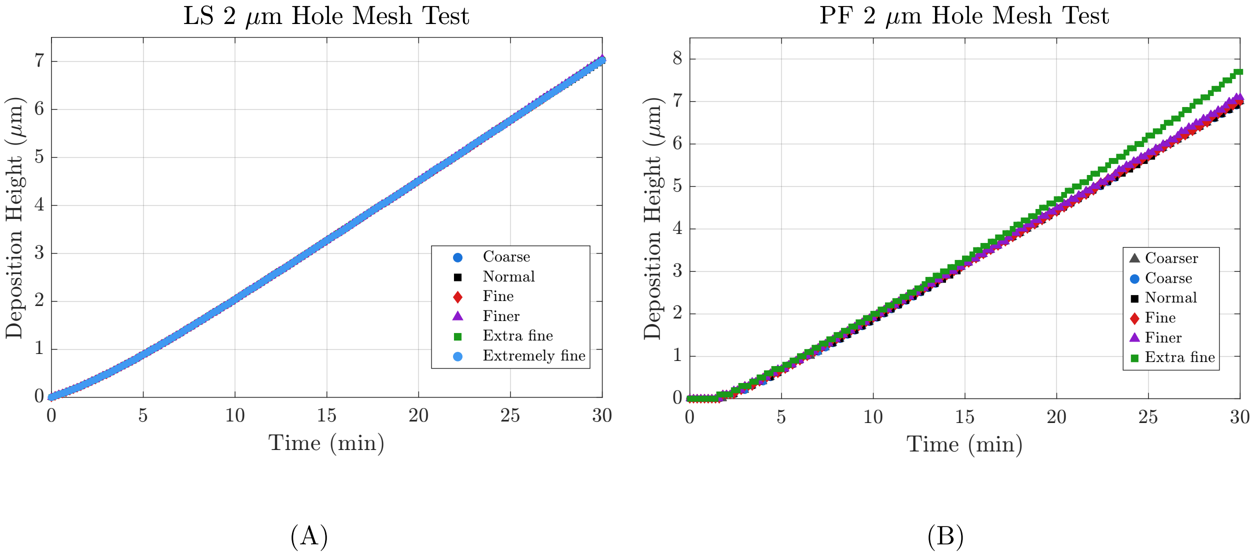

Figure S3: Mesh-convergence tests for a 2 μm hole geometry. (A) Level-set formulation. (B) Phase-field formulation. Progressively refined meshes produce nearly overlapping deposition-height curves over 30 min, supporting mesh-independent numerical predictions.

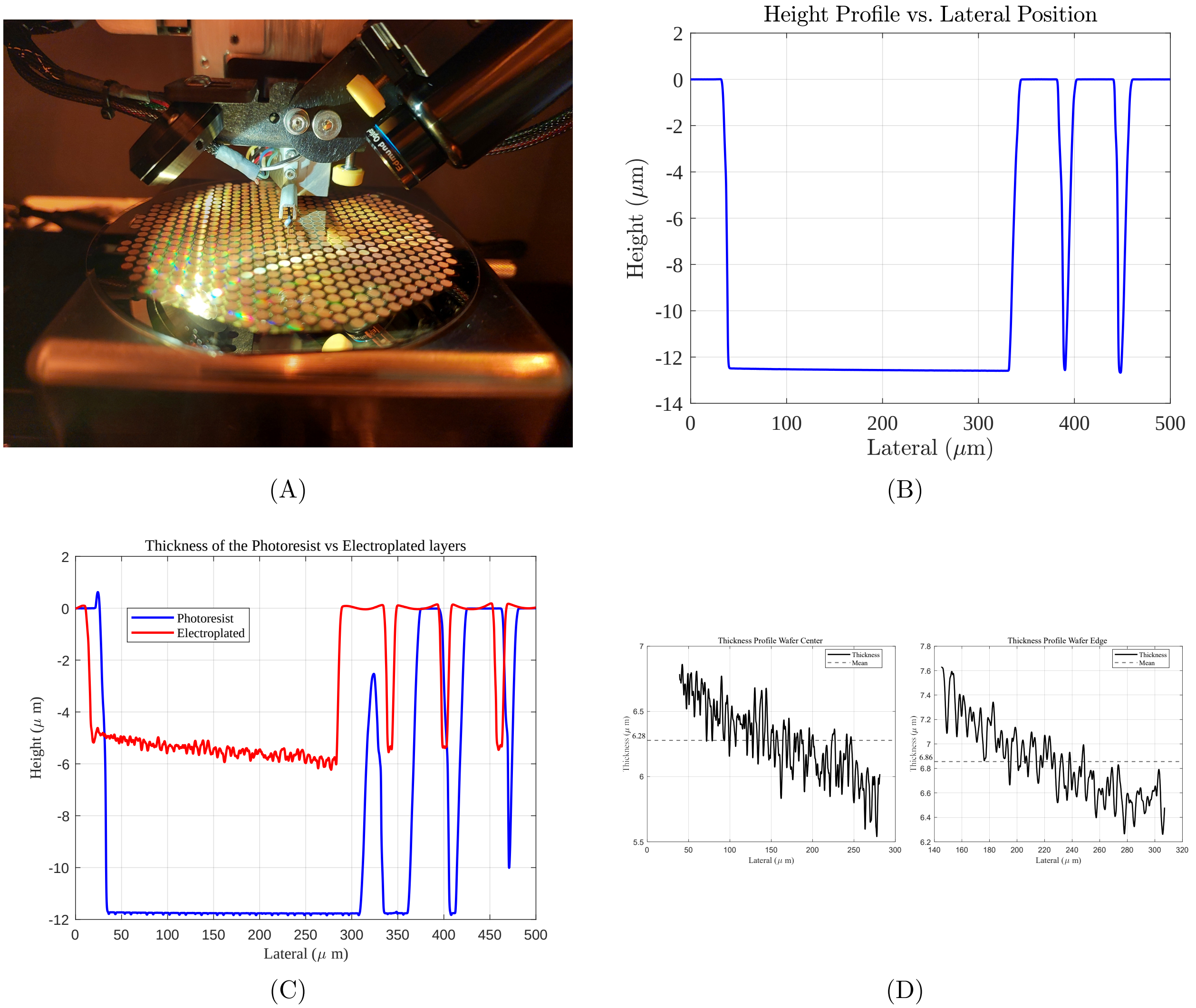

Figure S4: Profilometry-based validation of structural layer thickness. (A) Stylus profilometer setup on the patterned wafer. (B) AZ4562 pattern profile before plating, showing a resist height of approximately 12.5 µm. (C) Comparison of the pre-plating pattern profile with the electroplated structure. (D) Thickness measurements at the wafer center and edge, showing slightly higher deposition near the wafer edge and confirming final bar thickness in the tomography-compatible 6.5–8 µm range.

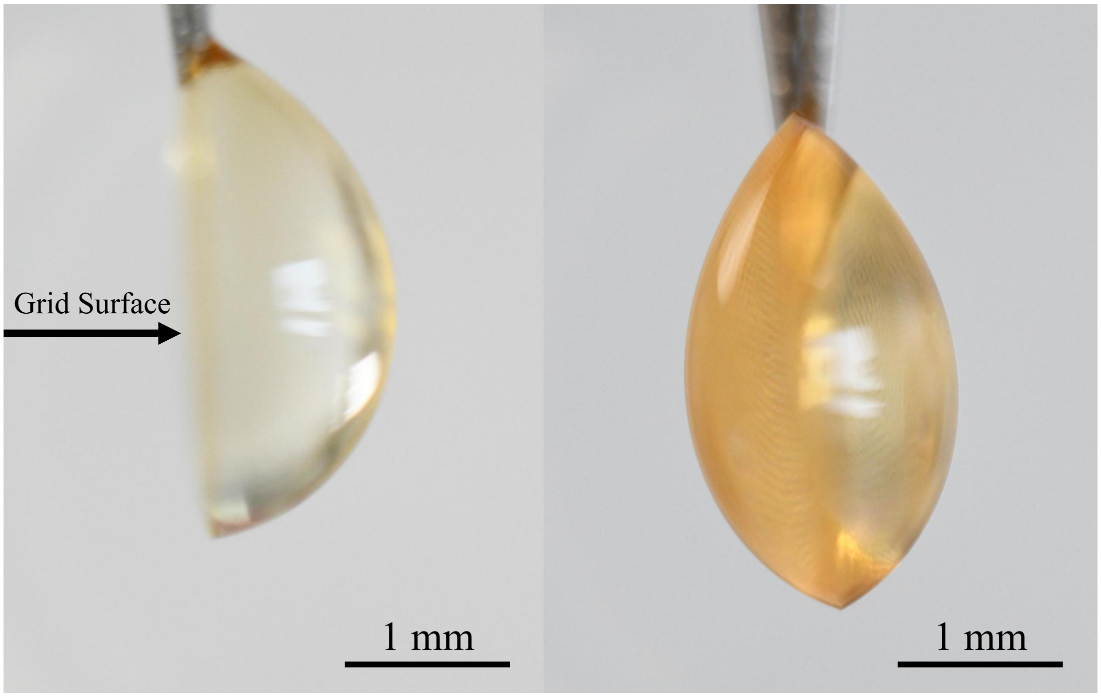

Figure S5: PEG-thiol wetting assay on all-gold grids. Left, unmodified grid; right, PEG-thiol-functionalized grid. The grid surface is oriented vertically and appears as the central line in each image. The PEG-thiol coating increases wettability, allowing the water droplet to pass through the foil and equilibrate on both sides.

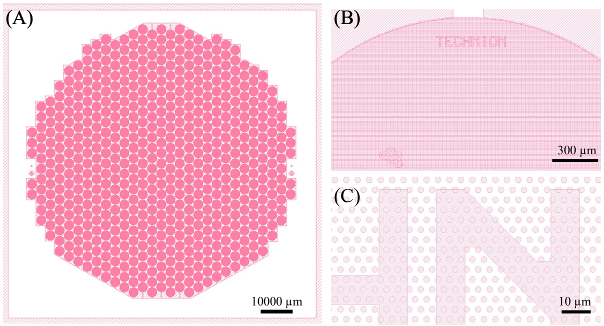

(A)

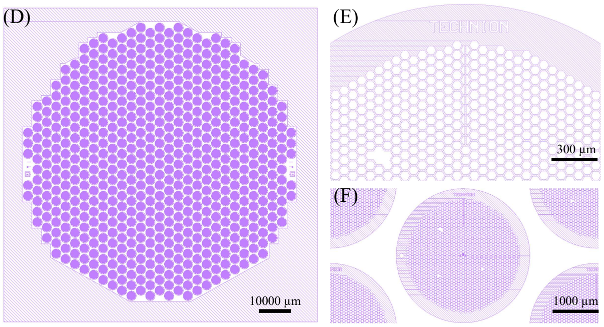

(B)

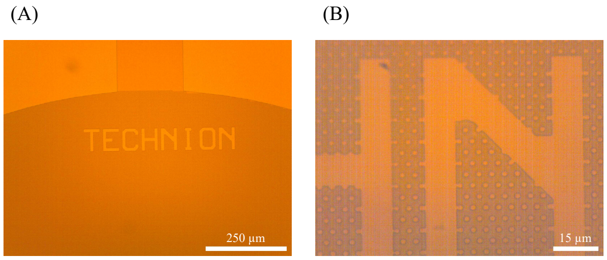

(C)

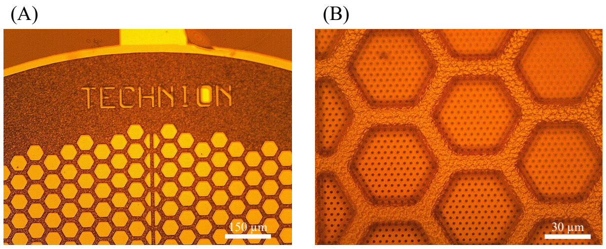

(D)

Figure S6: Circular-hole mask set and representative pattern-transfer steps. (A) Circular-hole foil-mask layout. (B) Mask layout used for the electroplated mesh. (C) Optical micrographs after exposure and development of the circular-hole foil pattern, including the grid perimeter, logo, and 2 µm hole array. (D) Optical micrographs after electroplating.

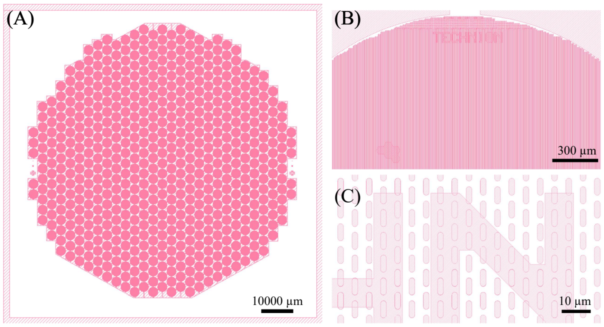

(A)

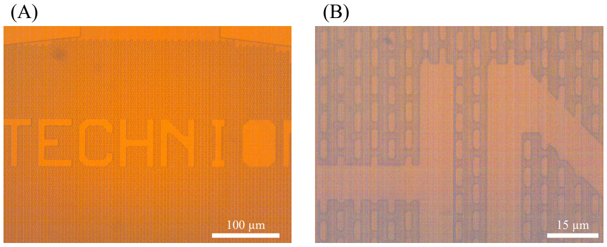

(B)

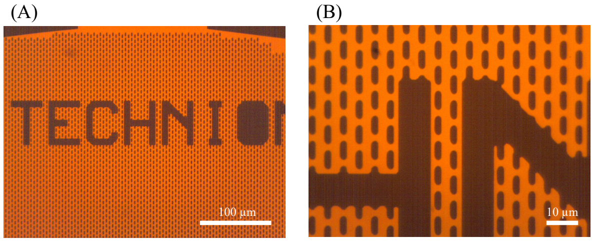

(C)

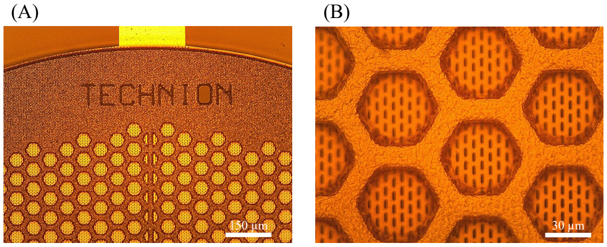

(D)

Figure S7: Oval-hole mask and representative fabrication sequence for anisotropic grids. (A) Layout of the oval-hole foil mask. (B) Oval-hole photoresist pattern after lithography. (C) Oval-hole foil after Ti/Au evaporation and lift-off. (D) Electroplated oval-hole grid after growth of the structural mesh.

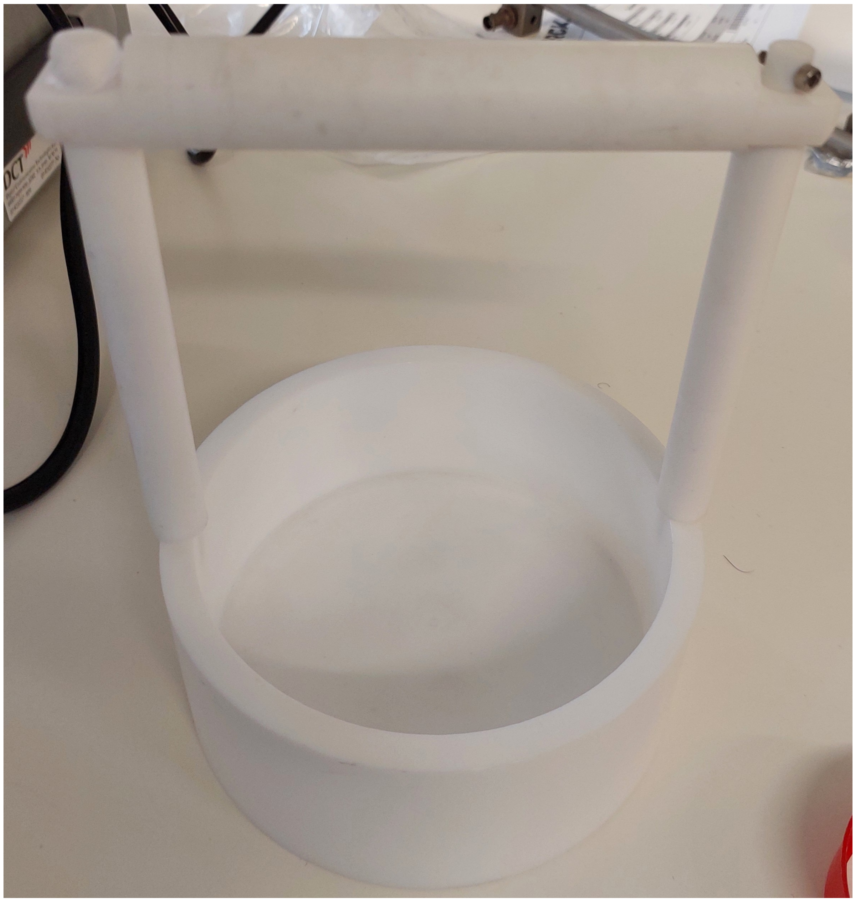

Figure S8: Custom HF immersion holder used during grid release. The Teflon fixture supports full-wafer immersion, drainage of the HF solution through side openings, controlled transfer between HF and deionized water, and reduced handling damage during release.

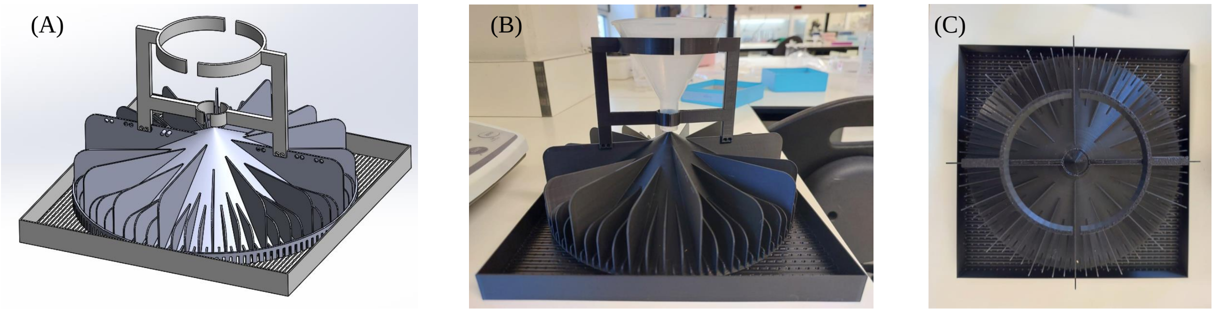

Figure S9: Grid retrieval and drying device. (A) CAD rendering. (B) Isometric view of the printed device. (C) Top view after use. Released grids are poured into the suspended funnel, partitioned into radial channels, and retained above a perforated lower plate that drains water while preserving the grids for drying and retrieval.

#### S3 Supplementary Tables

Table S1: Grid-level alignment-index statistics for oval-hole and circular-hole grids. Per-image AI values were first averaged within each grid to yield one grid-level value. Values are mean ± SD across grids ($n = 4$ grids per condition, from 2 independent experiments). Welch's $t$-test and Cliff's $\delta$ were calculated using grid-level means. Cliff's $\delta = 1.0$ indicates complete rank separation: every oval-hole grid exceeded every circular-hole grid with no overlap. For this complete-separation ranking, the exact Mann–Whitney $p$-value is 0.029.

| $h$ (deg) | Condition | AI mean ± SD | Welch $p$ | Cliff's $\delta$ |
|---|---|---|---|---|
| 10 | Oval | $0.1755 \pm 0.0138$ | $1.9 \times 10^{-3}$ | 1.00 |
| | Circular | $0.1206 \pm 0.0055$ | | |
| 20 | Oval | $0.3328 \pm 0.0234$ | $1.5 \times 10^{-3}$ | 1.00 |
| | Circular | $0.2387 \pm 0.0110$ | | |
| 30 | Oval | $0.4662 \pm 0.0285$ | $1.3 \times 10^{-3}$ | 1.00 |
| | Circular | $0.3533 \pm 0.0150$ | | |

Table S2: Sampling and preprocessing parameters used for actin-orientation analysis.

| Parameter | Value |
|---|---|
| Image-analysis software | Fiji/ImageJ Directionality plugin, local-gradient-orientation method |
| Independent experiments | 2 |
| Grids analyzed | 4 oval-hole grids and 4 circular-hole grids (2 per condition per experiment) |
| Images per grid | 8–11 images per oval-hole grid; 7–10 images per circular-hole grid |
| Total images | 38 oval-hole images, 36 circular-hole images (74 total) |
| Angular domain | Axial angles on $[0°, 180°)$; 90° defined as the long axis of the oval holes |
| Histogram bin width | Approximately 2° |
| Display smoothing | Circular Gaussian kernel, $\sigma = 6°$, area normalized; applied for display only |
| Per-image alignment metric | AI computed within half-window $h$ about 90° (Eq. S8) |
| Per-grid AI | Mean of per-image AI values within each grid |
| Primary analysis unit | Grid-level mean AI (4 grids per condition; images nested within grids, grids nested within experiments) |
| Group comparison | Welch two-sample $t$-test and Cliff's $\delta$ on per-grid means ($n = 4$ vs $n = 4$) |

Table S3: Axial anisotropy metrics for actin-orientation distributions computed over $[0^\circ, 180^\circ)$ from pooled group histograms.

| Metric | Value | Interpretation |
|---|---|---|
| $A_{\text{oval}}$ | $109.62 \times 10^{-5}$ | Mean-squared deviation from a uniform distribution for oval-hole grids |
| $A_{\text{circular}}$ | $2.08 \times 10^{-5}$ | Mean-squared deviation from a uniform distribution for circular-hole grids |
| $A_{\text{oval}}/A_{\text{circular}}$ | 52.52 | Relative increase in anisotropy for oval-hole grids |
| $\sqrt{A_{\text{oval}}}/\sqrt{A_{\text{circular}}}$ | 7.24 | Approximate fold increase in deviation magnitude for oval-hole grids |

Table S4: Electrochemical parameters used in the COMSOL electroplating model.

| Parameter | Value | Description |
|---|---|---|
| $C_{\mathrm{init}}$ | 60.9 mol $\mathrm{m}^{-3}$ | Initial electrolyte concentration |
| $T_0$ | 327 K | System temperature |
| $i_0$ | $2.8 \times 10^{-3}$ A $\mathrm{m}^{-2}$ | Exchange current density |
| $E_{\mathrm{eq,rel}}$ | 0 V | Relative equilibrium potential |
| $\phi_{s,\mathrm{anode}}$ | 0.15 V | Anode potential |
| $\phi_{s,\mathrm{cathode}}$ | $-0.15$ V | Cathode potential |
| $\alpha_c$ | 0.5 | Cathodic symmetry factor |
| $\alpha_a$ | 0.5 | Anodic symmetry factor |
| $z_{\mathrm{Au}}$ | 1 | Ionic charge, Au species |
| $z_{\mathrm{SO_3}}$ | $-2$ | Ionic charge, sulfite species |
| $D_{\mathrm{Au}}$ | $3.09 \times 10^{-9}$ $\mathrm{m}^2$ $\mathrm{s}^{-1}$ | Diffusivity of Au species |
| $D_{\mathrm{SO_3}}$ | $3.09 \times 10^{-9}$ $\mathrm{m}^2$ $\mathrm{s}^{-1}$ | Diffusivity of sulfite species |
| $M_{\mathrm{Au}}$ | 0.197 kg $\mathrm{mol}^{-1}$ | Molar mass of Au |
| $\rho_{\mathrm{Au}}$ | 19320 kg $\mathrm{m}^{-3}$ | Density of Au |

Table S5: Cryo-EM acquisition and processing parameters for the final apoferritin SPA dataset.

| Parameter | Value |
|---|---|
| Specimen | Apoferritin |
| Grid | Fabricated all-gold EM grid with 2 µm circular foil holes |
| Microscope | Thermo Fisher Glacios |
| Acceleration voltage | 200 kV |
| Detector | Falcon 4i direct electron detector |
| Energy filter | Selectris X |
| Energy-filter slit width | $\pm 5$ eV from zero-loss peak |
| Acquisition software | EPU |
| Nominal magnification | $x130,000$ |
| Pixel size | 0.89 Å |
| Total electron dose | 30 $e^{-}$/ Å$^{-2}$ |
| Defocus range | $-0.5$ to $-1.5$ µm |
| CTF fit-resolution cutoff | Better than 5.0 Å |
| Extraction box size | 300 pixels |
| Symmetry imposed | 0 |
| Initial particle images | 506,107 |
| Final particles images after 2D classification | 221,390 |
| Final map resolution | 2.15 Å |
| Resolution criterion | Gold-standard FSC = 0.143 |

### S4 Supporting Equations

For constant-current electroplating, the deposited thickness can be estimated from Faraday's law as

$$h(t) = \frac{j_c M_{\mathrm{Au}}}{z F \rho_{\mathrm{Au}}} \, t, \tag{S4}$$

where $j_c$ is the cathodic current density, $M_{\mathrm{Au}}$ is the molar mass of gold, $z$ is the number of electrons transferred per deposited Au ion, $F$ is Faraday's constant, and $\rho_{\mathrm{Au}}$ is the density of gold.

The geometric tilt-access limit follows from the lateral displacement of a central electron ray,

$$\Delta x = t \tan\theta, \tag{S5}$$

and the requirement $\Delta x < L$ with $L = W/2$ for a hexagonal window of across-flats width $W$. This gives

$$t_{\max} = \frac{L}{\tan\theta}, \qquad \theta_{\max} = \arctan\left(\frac{L}{t}\right). \tag{S6}$$

For display of actin-orientation histograms, circular Gaussian smoothing was applied as

$$y_i^{(\mathrm{sm})} = \sum_{k=-r}^{r} g_k \, y_{(i-k) \bmod N}, \qquad g_k = \frac{1}{Z} \exp\left[-\frac{1}{2}\left(\frac{k}{\sigma_{\mathrm{bins}}}\right)^2\right], \tag{S7}$$

where $y_i$ is the unsmoothed histogram, $N$ is the number of bins, $g_k$ are normalized Gaussian weights with $\sum g_k = 1$, and the modulo operation enforces circular continuity over $[0, 180)$.

The alignment index used for the cell-orientation analysis is

$$\mathrm{AI} = \frac{\sum_{|\theta - 90^\circ| \le h} w(\theta)}{\sum_{\theta \in [0,180)} w(\theta)}, \tag{S8}$$

where $w(\theta)$ is the gradient-weighted orientation count and $h$ is the half-window around the substrate axis.

The axial anisotropy metric is

$$A = \sum_{i=1}^{N} \left(P(\theta_i) - \frac{1}{N}\right)^2, \tag{S9}$$

where $P(\theta_i)$ is the normalized probability in angular bin $i$ and $\sum_i P(\theta_i) = 1$.
